# Pressure for rapid and accurate mate recognition promotes avian-perceived plumage sexual dichromatism in true thrushes (genus: *Turdus*)

**DOI:** 10.1101/2021.08.12.456147

**Authors:** Alec B. Luro, Mark E. Hauber

## Abstract

Ecological conditions limiting the time to find a compatible mate or increasing the difficulty in doing so likely promote the evolution of traits used for species and mate recognition. Conspicuous traits that signal an individual’s species, sex, and breeding status reduce the challenge of identifying a compatible conspecific mate, and should be more common in migratory rather than sedentary species, species with shorter breeding seasons, and species breeding under high sympatry with many closely-related heterospecifics. Here, we tested this recognition hypothesis for promoting plumage sexual dichromatism in the true thrushes (*Turdus* spp.), a large and diverse genus of passerine birds. We used receptor-noise limited models of avian vision to quantify avian-perceived chromatic and achromatic visual contrasts between male and female plumage patches and tested the influence of breeding season length, spatial distribution, and sympatry with other *Turdus* species on plumage dichromatism. As predicted, we found that 1) true thrush species with migratory behaviour have greater plumage sexual dichromatism than non-migratory species, 2) species with longer breeding seasons have less plumage sexual dichromatism, and 3) greater numbers of *Turdus* thrush species breeding in sympatry is associated with more plumage sexual dichromatism. These results suggest that social recognition systems, including species and mate recognition, play a prominent role in the evolution of plumage sexual dichromatism in true thrushes.

## Introduction

Species recognition is necessary in sexually reproducing lineages for individuals to find compatible mates and produce viable offspring [1,2]. Conspicuous traits signaling species and sex identity increase the ease and speed of mate recognition by reducing the effort, error, and time involved when searching for compatible mates and lessen the likelihood of mating with heterospecifics [3]. Traits used in species and mate recognition may also serve as signals of status to conspecifics and reduce costly conflicts over resources and mates [4]. Accordingly, distinct traits facilitating mate recognition should be more likely to arise and be maintained under conditions that increase both the difficulty of finding a compatible mate and degree of resource competition among conspecifics and closely-related species. Conditions likely to favour traits signaling individuals’ species, sex, and breeding status include higher sympatry with many closelyrelated species, limited time to find compatible breeding mates, and lower rates of encounter with potential breeding mates [1].

In birds, plumage colour is a highly conspicuous trait signaling species and (often) sex identity [5,6]. Plumage sexual dichromatism, or the distinct set of differences in the appearance of male and female feather colours and patterns, is common in birds and is usually attributed to different natural and sexual selection pressures on males and females [7–11]. Plumage sexual dichromatism results in a visibly perceivable trait useful for recognizing an individual’s species, sex, and breeding status (e.g., in species with sex-specific delayed plumage maturation, see [12]), reducing the time and effort expended to identify a suitable mate [13,14]. Evidence in favour of this recognition hypothesis for sexual dichromatism in birds includes a positive association of greater plumage sexual dichromatism with migratory behaviour and shorter breeding seasons [9], both of which reduce the amount of time available to search and find suitable mates and successfully breed. Additional support for the recognition hypothesis includes a consistent pattern of greater plumage sexual dichromatism and plumage colour elaboration in avian species that reside on mainland continents and have large geographic ranges in comparison to species that do not migrate, reside on islands, and have limited breeding ranges [10,15–23].

Moreover, plumage sexual dichromatism likely plays a role in hybridization avoidance via reproductive character displacement to facilitate species and mate recognition, especially among closely-related species. For example, in *Ficedula* flycatchers, female choice selects for divergent male plumage colouration across populations and species, leading to male character displacement and reduced rates of interspecific hybridization [24–26]. More broadly and across taxa, greater plumage dichromatism is positively associated with higher breeding sympatry with closely-related heterospecifics. Among a large sample of passerine sister species pairs, transitions from allopatry to parapatry and increases in geographic range overlaps are positively correlated with greater plumage dichromatism [27]. Greater plumage sexual dichromatism has also been found to be positively associated with greater avian species divergence and richness [28,29]. Among passerine sister species pairs, more pronounced changes in male rather than female plumage colouration in sexually-dichromatic species suggest that female choice and male-male competition often lead to concurrent increases in sexual dichromatism and speciation events [28]. Therefore, plumage sexual dichromatism may be a selected trait for facilitating species and mate recognition when closely-related species have sympatric breeding ranges [5,30].

True thrushes (*Turdus* spp.) are an exceptionally diverse monophyletic genus of passerine birds consisting of about ~86 species distributed across the globe (Fig. 1). The true thrushes are an ideal passerine clade for examining the recognition hypothesis for plumage sexual dichromatism because plumage sexual dichromatism and migratory behaviours vary substantially between species and sexual dichromatism has evolved multiple times in thrushes across the world [31,32]. Hybridization also occurs in some, but not all, *Turdus* species, indicating that some sympatric *Turdus* species can successfully interbreed. A particulary well-documented example of hybridization in true thrushes occurs at large hybrid zone between four *Turdus* species (*T. atrogularis*, *T. eunomus*, *T. naumanni*, *T. ruficollis*) in north-central Asia [33]. Further, plumage sexual dichromatism in true thrushes often coincides with age and breeding status in male thrushes. Delayed plumage maturation in males is common among true thrushes [34–36], where males have “female-like” plumage colouration during their first breeding season and develop typical breeding adult male plumage for subsequent breeding seasons. The presence of delayed plumage maturation and distinct juvenal plumage may serve as a signal of a young male’s sexual immaturity in order to reduce levels conspecific aggression from older adults [36] and also suggests that female thrushes prefer older males with prominent adult plumage as breeding mates.

**Figure 1:**
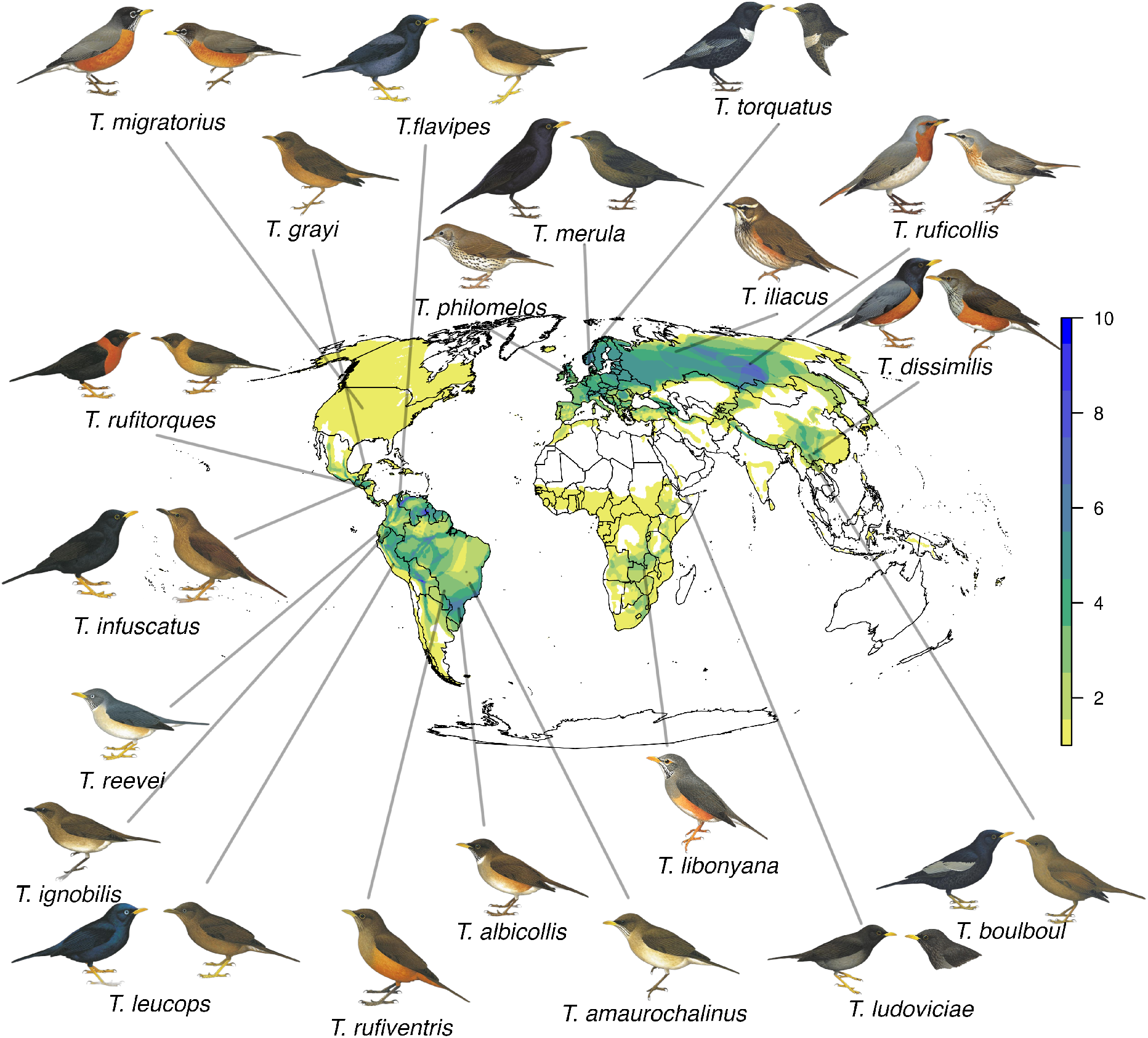
Breeding ranges of all recognized *Turdus* species from BirdLife International, with representative species’ males and females shown for species with plumage sexual dichromatism. The color scale indicates the number of *Turdus* thrush species in sympatry with overlapping breeding ranges. Illustrations used with permission from HBW Alive/Lynx Edicions

Overall, ecological conditions that increase the time and degree of difficulty in finding a suitable conspecific mate should select for phenotypic traits that reliably signal species and sex identity. Across various bird lineages, greater plumage dichromatism is present in species that are i) migratory rather than nonmigratory, ii) have shorter breeding seasons, iii) live on mainlands rather than islands, iv) have larger breeding ranges (distributions), and v) breed in sympatry with more closely-related species. These patterns suggest that ecological circumstances where rapid and accurate mate recognition is challenging strongly favour the evolution and maintenance of prominent plumage sexual dichromatism in birds. Here, we test these predictions of the recognition hypothesis for plumage sexual dichromatism by evaluating the potential influences of breeding timing, spacing, and sympatry on plumage dichromatism in *Turdus* thrushes (Fig. 2).

**Figure 2:**
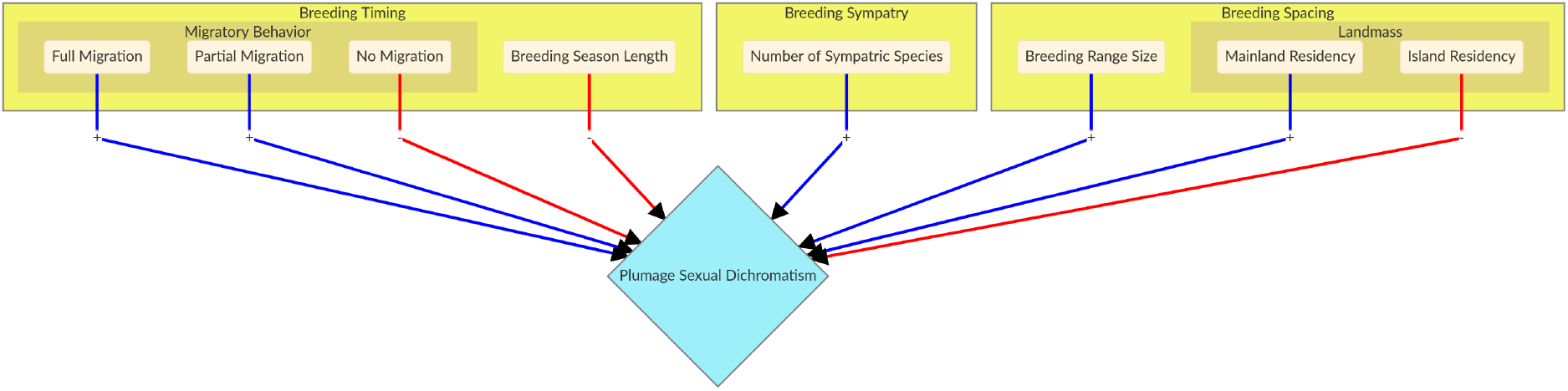
Hypotheses and predictions for each model (large yellow boxes). Arrow colours indicate predicted correlation, positive (blue) and negative (red)

## Methods

Initial pre-registration of the study’s methods and analyses are available on Open Science Framework [37].

### Plumage sexual dichromatism

A total of N=77 *Turdus* thrush species (approximately ~89% of all known true thrush species) were sampled for plumage spectral reflectance using prepared bird skin specimens at the American Museum of Natural History in New York City and the Field Museum in Chicago, USA. Reflectance measurements spanning 300-700nm were taken in triplicate from the belly, breast, throat, crown, and mantle plumage patches [38] of each individual. N=3 male and N=3 female individuals were measured for most species (exceptions: *T. lawrencii*, N=2 males and N=2 females; *T. swalesi*, N=1 male and N=1 female). Reflectance spectra were measured using a 400 μm fiber optic reflection probe fitted with a rubber stopper to maintain a consistent measuring distance of 3 mm and area of 2 mm2 at a 90° angle to the surface of the feather patch. Measurements were taken using a JAZ spectrometer with a pulsed-xenon light source (Ocean Optics, Dunedin, USA) and we used a diffuse 99% reflectance white standard (Spectralon WS-1-SL, Labsphere, North Sutton NH, USA).

We applied a receptor-noise limited visual model [39] of the European Blackbird (*T. merula*) visual system [40] in the *pavo* [41] package in R v4.0.0 [42] to calculate avian-perceived chromatic and achromatic visual contrast (in units of “Just-Noticeable Differences”,or JNDs) of male vs. female plumage patches for all sampled *Turdus* species. Chromatic and achromatic JNDs were calculated for male-female pairs within each species (i.e., N=9 JND values calculated per patch for each species where N=3 males and N=3 females sampled), and then JND values were averaged for each species’ respective plumage patches. Under ideal laboratory conditions, 1 JND is generally considered to be the discriminable threshold past which an observer is predicted to be able to perceive the two colours as different. However, natural light environments vary both spatially and temporally [43], bringing into question the accuracy of a 1 JND threshold for generalizing visual contrast under natural conditions. Therefore, we calculated the total number of sexually-dichromatic plumage patches per species (out of N=5 measured patches) as the number of plumage patches with average JND values > 1, 2, or 3 to account for uncertainty in visual discrimination thresholds due to variation in psychophysical and ambient lighting conditions affecting the strength of between-sex plumage visual contrast [44]. Additionally, we modeled the number of divergent plumage patches (at the three different JND thresholds listed above) within sexes and between different sympatric species under different levels of breeding range overlap (10% increments between 0-90%; Fig. S1).

### Life History Data

#### Breeding Timing Model

We collected data on migration behaviour and breeding season length from *Thrushes* [31] and the *Handbook of the Birds of the World* [45]. We assigned three different kinds of migratory behaviour: 1) *full migration* when a species description clearly stated that a species “migrates”, 2) *partial migration* when a species was described to have “altitudinal migration”, “latitudinal migration” or “movement during non-breeding season”, or 3) *sedentary* when a species was described as “resident” or “sedentary”. Breeding season length was defined as the number of months the species breeds each year.

#### Breeding Sympatry Model

Species’ breeding ranges were acquired from *BirdLife International* [46]. We calculated congener breeding range overlaps (as percentages) using the *letsR* package in R [47]. We then calculated the number of sympatric species as the number of congeners with breeding ranges that overlap >30% with the focal species’ breeding range [27]. Comparisons of the number of sexually-dimorphic plumage patches vs. the number of sympatric species among different breeding range overlap thresholds are provided in Supplementary Figure 2.

#### Breeding Spacing Model

Species’ breeding range sizes (in km2) were acquired using the *BirdLife International* breeding range maps. Species’ island vs. mainland residence was also determined using breeding ranges from *BirdLife International*. Mainland residence was assigned if the species had a breeding range on any continent and Japan. Island residence was assigned to species having a breeding range limited to a non-continental landmass entirely surrounded by a marine body of water.

#### Statistical modeling

We used phylogenetically-corrected Bayesian multilevel logistic regression models using the *brms* v2.13.0 package [48] in R v4.0.0 [42]. We modeled plumage sexual dichromatism responses as the number of sexually-dichromatic patches > 1, 2, or 3 chromatic and achromatic JNDs. Plumage dichromatism responses were modeled as binomial trials (N=5 plumage patch “trials”) to test for associations with breeding timing, breeding sympatry and breeding spacing. For all phylogenetically-corrected models, we used the *Turdus* molecular phylogeny from Nylander et al. (2008) [49] to create a covariance matrix of species’ phylogenetic relationships. All models used a dataset of N=67 out of the *Turdus* species for which all the types of data (see above) were available.

Our breeding timing models included the following predictors: z-scores of breeding season length (mean-centered by *μ* = 5.4 months, and scaled by one standard deviation *σ* = 2.3 months), migratory behaviour (no migration as the reference category versus partial or full migration), and their interaction. *Breeding sympatry* models included the number of sympatric species with greater than 30% breeding range overlap as the only predictor of the probability of having a sexually-dichromatic plumage patch. *Breeding spacing* models included transformed breeding range size (km2) and breeding landmass (mainland as the reference category versus island). We also ran null models (intercept only) for all responses. All models’ intercepts and response standard deviations were assigned a weakly informative prior (Student T: df = 3, location = 0, scale = 10) [50], and predictor coefficients were assigned flat uninformative priors. We ran each model for 6,000 iterations across 6 chains and assessed Markov Chain Monte Carlo (MCMC) convergence using the Gelman-Rubin diagnostic (Rhat) [50]. We then performed k-fold cross-validation [51] to assess each model’s accuracy in predicting plumage sexual dichromatism of randomly-selected samples of *Turdus* thrush species, refitting each model *K*=16 times. For each k-fold, the training dataset included a randomly selected set of 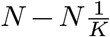 or N≈63 species, and the testing dataset included 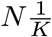 or N≈4 species not included in the training dataset. Finally, we compared differences between the models’ expected log pointwise predictive densities (ELPD) to assess which model(s) best predicted the probability of having a sexually-dichromatic plumage patch. [51].

Models’ predictor effects were assessed using 90% highest-density intervals of the posterior distributions and probability of direction, the proportion of the posterior distribution that shares the same sign (positive or negative) as the posterior median [52], to provide estimates of the probability of that a predictor has an entirely positive or negative effect on the presence of sexually-dimorphic plumage patches. We assume predictor estimates with a probability of direction ≥ 0.90 to be indicative of a reliable existence of a predictor’s effect on sexually-dimorphic plumage patches [52].

## Results

### Avian visual modeling

Among N=77 *Turdus* species, the following proportion have sexually monomorphic plumage (combined achromatic and chromatic JND thresholds): 1.3% (n=1 species) have no sexually-dimorphic patches > 1 JND, 44% (n=34 species) have no dimorphic patches > 2 JND, and 63% (n=49 species) have no dimorphic patches > 3 JND (Table S1). Additional proportions of *Turdus* species with sexually-dimorphic achromatic or chromatic plumage patches are available in Table S2. When comparing within sexes between sympatric species (i.e., following [27] at least a 30% overlap in breeding ranges: n=39 species with at least one sympatric species and a median of n=6 sympatric species per focal species), the median number of avian-discriminable plumage patches between species is 1 or greater for all three achromatic and chromatic JND thresholds except for sympatric females at a chromatic JND threshold > 3 (Fig. S1).

### Model comparisons

*Breeding sympatry*, *breeding timing*, and *breeding spacing* performed considerably better than *intercept-only* (null models) in predicting the probability of a species having a sexually-dimorphic plumage patch. We obtained N ≥ 4000 effective posterior samples for each model parameter and all models’ Markov Chains (MCMC) successfully converged (Rhat = 1 for all models’ parameters). All *breeding sympatry*, *breeding timing*, and *breeding spacing* models performed similarly well and substantially better than *intercept only* models in predicting the probability of having a sexually-dimorphic plumage patch with achromatic JND values > 1, 2, or 3 (Table 1; all models predicting achromatic plumage patches had ELPD values within 4, following the convention of [53]). Among models predicting the probability of having a sexually-dichromatic plumage patch with chromatic JND values >1, 2, or 3, all *breeding sympatry*, *breeding timing*, and *breeding spacing* models performed much better than *intercept only* models, and *breeding sympatry* models had the top predictive performance (Table 1; *breeding sympatry* models all have ELPD =0, only the *breeding spacing* models predicting dichromatic plumage patches had similar predictive performance).

**Table 1:** Expected log pointwise predictive densities (ELPD) differences and kfold information criterion values of models (ELPD Difference ± standard error (kfold IC ± standard error)). Values closest to zero indicate greater model prediction performance.

|  |  | Model |  |  |  |
| --- | --- | --- | --- | --- | --- |
| Plumage Metric | JND Threshold | Breeding Sympatry | Breeding Timing | Breeding Spacing | Intercept Only |
| Achromatic |  |  |  |  |  |
|  | 1 JND | 0 ± 0 (-122.17 ± 0.67) | -2.51 ± 2.49 (-124.68 ± 2.38) | -2.59 ± 1.01 (-124.76 ± 1.04) | -21.69 ± 7.36 (-143.87 ± 7.51) |
|  | 2 JND | 0 ± 0 (-98.94 ± 7.56) | -1.19 ± 3.95 (-100.13 ± 9.22) | -0.7 ± 1.34 (-99.64 ± 7.92) | -52.42 ± 12.67 (-151.36 ± 13.4) |
|  | 3 JND | -0.04 ± 1.4 (-85.4 ± 8.91) | -1.7 ± 4.41 (-87.07 ± 10.71) | 0 ± 0 (-85.37 ± 8.76) | -28.54 ± 10.02 (-113.91 ± 13.65) |
| Chromatic |  |  |  |  |  |
|  | 1 JND | 0 ± 0 (-115.75 ± 2.95) | -5.67 ± 3.55 (-121.42 ± 2.28) | -2.73 ± 3.4 (-118.49 ± 2.67) | -14.8 ± 7.22 (-130.55 ± 7.05) |
|  | 2 JND | 0 ± 0 (-88.47 ± 8.77) | -3.8 ± 4.46 (-92.27 ± 10.01) | -3.32 ± 5.29 (-91.79 ± 10.91) | -50.53 ± 14.49 (-139 ± 16.77) |
|  | 3 JND | 0 ± 0 (-62.77 ± 10.41) | -8 ± 4.32 (-70.77 ± 12.29) | -4.43 ± 3.9 (-67.2 ± 11.72) | -47.63 ± 15.34 (-110.4 ± 20.01) |

### Achromatic plumage sexual dichromatism predictors

Migratory behaviour and shorter breeding season lengths were strongly associated with greater odds of a species having achromatic plumage sexual dichromatism. All model predictors’ effect estimates are provided as the posterior median odds-ratio (OR) and 90% highest-density interval (HDI) in Table 2. Among predictors of achromatic sexually-dimorphic plumage patches, only predictors included in the *breeding timing* model have predictors with probability of direction (*pd*) values ≥ 0.90 (Table 2). Specifically, longer breeding season length was associated with lower odds of a species having a sexually-dimorphic plumage patch with achromatic JND > 2 (breeding season length, OR [90% HDI] = 0.10 [0.01, 1.1], 89.5% decrease in odds per 2.3-month increase in breeding season) and JND > 3 (breeding season length, OR [90% HDI] = 0.25 [0.03, 1.5], 75% decrease in odds per 2.3-month increase in breeding season). Additionally, full migratory behaviour, rather than no migratory behaviour, was associated with greater odds of a species having a sexually-dimorphic plumage patch with achromatic JND > 1 (full migration, OR [90% HDI] = 4.97 [0.95, 24.4]), JND > 2 (full migration, OR [90% HDI] = 66.5 [3.2, 1802.4]) and JND > 3 (OR [90% HDI] = 22.3 [1.6, 307.9]). Finally, both full and partial migratory behaviour, rather than no migration behaviour, in conjunction with longer breeding season lengths are associated with greater odds of a species having a sexually-dimorphic plumage patch with achromatic JND > 1 (breeding season length x full migration, OR [90% HDI] = 4.84 [0.67, 39.6]), JND > 2 (breeding season length x full migration, OR = 66.3 [0.59, 11415.7]; breeding season length x partial migration, OR [90% HDI] = 20.7 [0.9, 589.1]) and JND > 3 (breeding season length x partial migration, OR [90% HDI] = 8.28 [0.76, 109.1]).

**Table 2:**
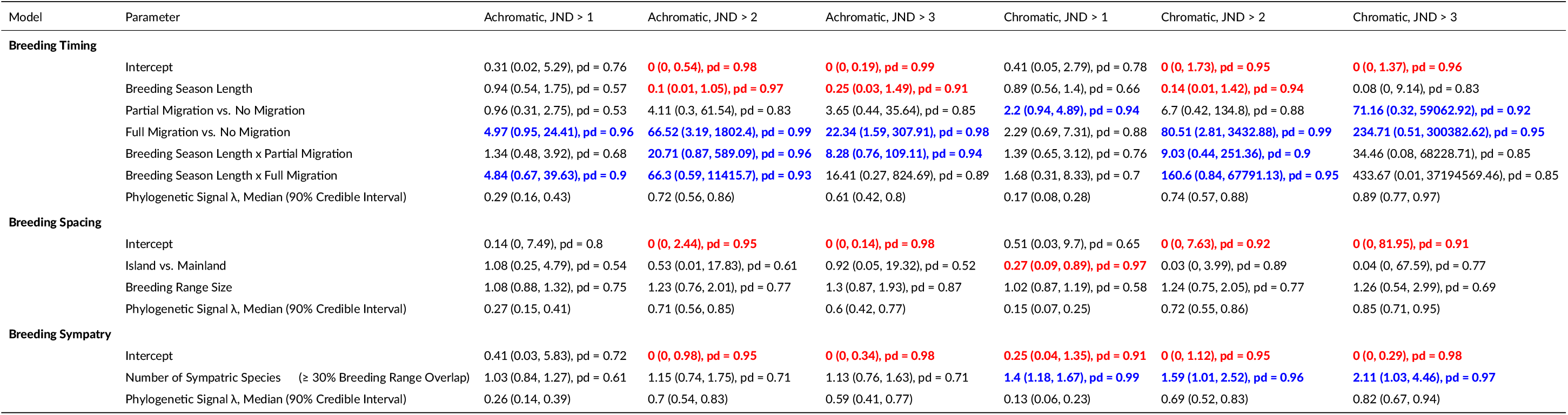
Model predictor effect estimates (posterior median odds ratio and 90% highest-density interval) on the presence of a plumage patch with achromatic or chromatic visual contrast values >1, 2, and 3 JND. Model effects with a probability of direction (pd) value ≥0.90 are bolded in red for a negative effect and blue for a positive effect on plumage dichromatism. Phylogenetic signal (λ) for each model is provided as the median and 90% credible interval of the intraclass correlation coefficient among species.

### Chromatic plumage sexual dichromatism predictors

Migratory behaviour, shorter breeding season lengths, and larger numbers of sympatric *Turdus* species were strongly associated with greater odds of a species having chromatic plumage sexual dichromatism. Among predictors of *breeding timing* models predicting chromatic sexually-dimorphic plumage patches, longer breeding season length was associated with lower odds of a species having a plumage patch with chromatic JND > 2 (OR [90% HDI] = 0.14 [0.01, 1.42], 86% reduction in odds per 2.3 month increase in breeding season). Both full and partial migratory behaviour rather than no migration are associated with greater odds of a species having a plumage patch JND > 1 (partial migration, OR [90% HDI] = 2.2 [0.94, 4.9]), JND > 2 (full migration, OR [90% HDI] = 80.51 [2.8, 3432.9]) and JND > 3 (partial migration, OR [90% HDI] = 71.2 [0.32, 59062.9]; full migration, OR [90% HDI] = 234.7 [ 0.51, 300382.6]). For *breeding spacing models*, island residency rather than mainland residency was associated with lower odds of having a plumage patch > 1 chromatic JND (island, OR [90% HDI] = 0.27 [0.09, 0.89]). Finally, more *Turdus* species in sympatry was associated with higher odds of a species having a sexually-dimorphic chromatic plumage patch with JND > 1 (number of sympatric species, OR [90% HDI] = 1.4 [1.18, 1.67], 40% increase in odds per each additional sympatric species), JND > 2 (sympatric species, OR [90% HDI] = [1.01, 2.52], 59% increase in odds per each additional sympatric species), and JND > 3 (sympatric species, OR [90% HDI] = 2.11 [1.03, 4.46], 111% increase in odds per each additional sympatric species).

## Discussion

Our results provide comparative correlative evidence in support of predictions of the recognition hypothesis for plumage sexual dichromatism in true thrushes. We used a receptor-noise limited model of *Turdus merula* vision [39,40] to measure avian-perceivable visual contrast of plumage colours and found that the odds of plumage sexual dichromatism are much greater for *Turdus* thrush species that have full or partial migration rather than no migration, have relatively short breeding seasons, and are in sympatry with many other true thrush species (Table 1,2). Our results align with prior comparative studies of avian plumage sexual dichromatism where strong associations of sexual dichromatism with greater migratory behaviour [10] and more sympatric taxa [27] were found among many species of different passerine families.

Further, we determined that sympatric *Turdus* species have distinguishable plumage colouration differences from one another when measuring plumage appearance from the avian visual perspective (Fig. S1). Divergent plumage colouration within sexes between closely-related species indicates that plumage sexual dichromatism may have evolved to facilitate species and mate recognition in *Turdus* species breeding under higher sympatry with other true thrushes. However, we cannot directly determine if the plumage sexual dichromatism in sympatric *Turdus* species is the result of reproductive character displacement. We do not know if past changes in species’ plumage sexual dichromatism occurred before or during periods of sympatry with other *Turdus* species. Regardless, present-day plumage sexual dichromatism and perceivable differences in plumage colouration between sympatric species likely reduces the challenge of finding compatible mates by signaling an individual’s sex, breeding status, and species. For example, the four species *Turdus* hybrid zone in north-central Asia [33] is a particularly striking example where reproductive character displacement has likely occurred and all four species exhibit strong plumage sexual dichromatism (Fig. S3). Comparing within sexes between sister species pairs of *T.ruficollis* and *T.atrogularis*, and *T.naumanni* and *T.eunomus* [49], plumage patterns of the species pairs are nearly identical except for a divergence in colour. *T.ruficollis* and *T.atrogularis* share similar facial and throat colouring patterns, with the main difference being red colouration in *T.ruficollis* in opposition to the black colouration of *T.atrogularis*. In the second species pair, *T.naumanni* has red ventral plumage colouration and *T.eunomus* has black ventral plumage colouration.

Previous studies have found that closely-related sympatric species tend to have more similar plumage appearance than expected if plumage colouration patterns had evolved to facilitate species recognition via reproductive character displacement [54,55]. The potential lack of major plumage colour divergence among closely-related sympatric species may be attributable to constraints imposed by a shared light environment on colour signal efficiency [56], or similar natural selection pressures (e.g., predators, parasites, and weather). Generally, despite greater similarity in plumage appearance in comparison to allopatric species, closely-related sympatric species can still have substantially different and biologically-relevant differences in achromatic or chromatic interspecific visual contrast of plumage patches when measuring plumage colouration differences from the avian visual perspective (as we have found in our analyses).

## Conclusions

Patterns of plumage sexual dichromatism in true thrushes (*Turdus*) are consistent with select predictions of the recognition hypothesis for plumage sexual dichromatism. Migratory behaviour and limited breeding seasons reduce the amount of time available to find a mate, and greater plumage sexual dichromatism may help migratory species find compatible mates more rapidly. Greater plumage sexual dichromatism in *Turdus* species under sympatry with other true thrush species also supports the possibility that increased plumage sexual dichromatism may be the result of reproductive character displacement. Therefore, greater plumage sexual dichromatism likely increases the speed and accuracy of finding a compatible breeding mate, reduces species and mate recognition errors, and decreases hybridization.

## Acknowledgements

We thank the American Museum of Natural History in New York City and Field Museum of Chicago for access to museum specimens used in this study. We also thank the Department of Evolution, Ecology, and Behavior at the University of Illinois for funding and support. MEH was funded by the University of Illinois Harley Jones Van Cleave Professorship. We are grateful for the extensive feedback and comments from Becky Fuller, Jeffrey Hoover, and Al Roca that greatly improved the manuscript.

## Data Accessibility

Data and code used for the analyses can be found at https://github.com/aluro2/Turdus-Dichromatism.

## Author Contributions

**Alec Luro:** Conceptualization, Investigation, Methodology, Software, Formal Analysis, Data Curation, Visualization, Writing-Original Draft, Writing-Review & Editing. **Mark Hauber:** Conceptualization, Resources, Supervision, Project administration, Funding acquisition, Writing-Review & Editing.

## Supplementary Tables and Figures

**Table S1:**
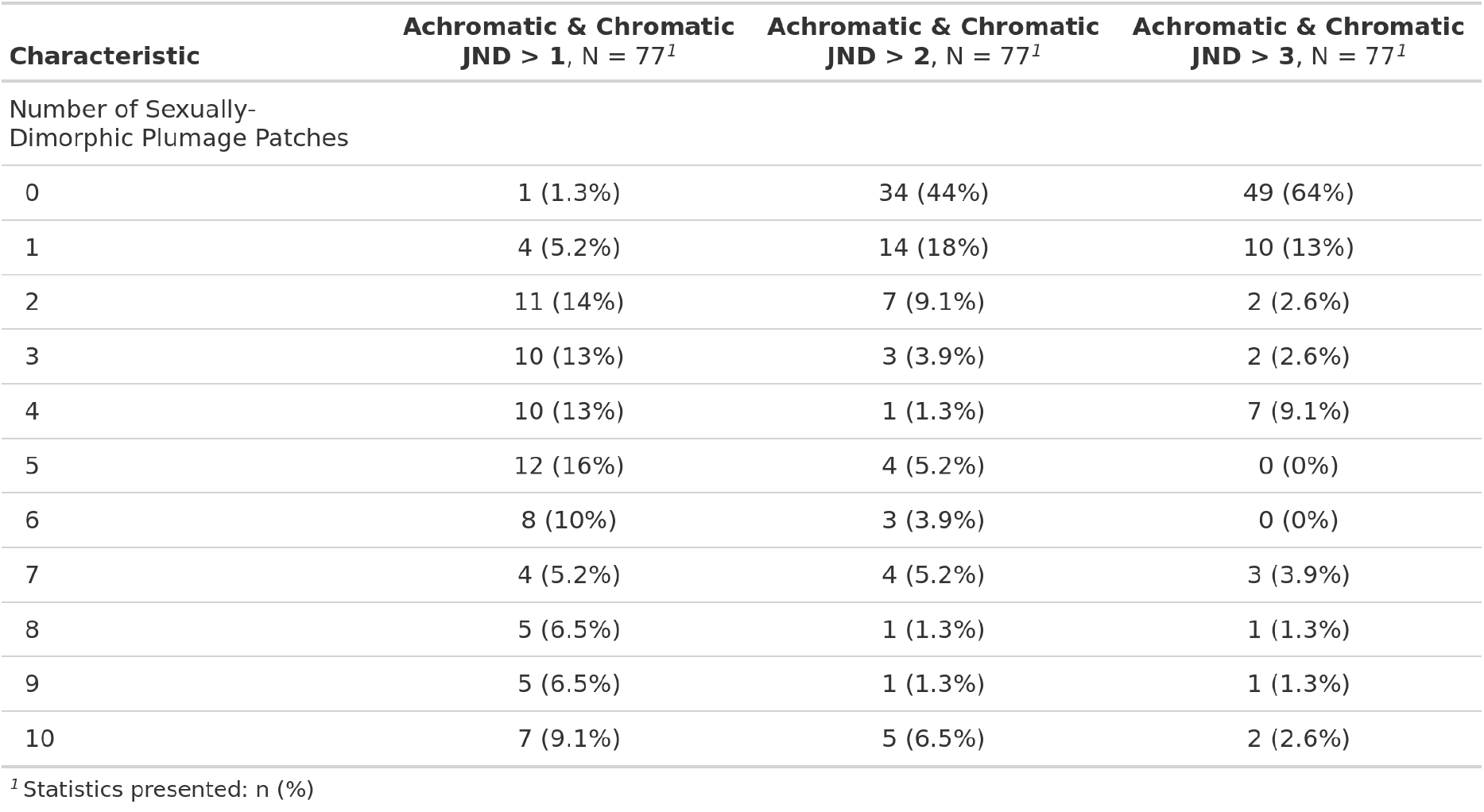
Number of sexually-dimorphic plumage patches for combined achromatic and chromatic just noticeable differences (JND) thresholds by number of *Turdus* thrush species (% of species).

**Table S2:**
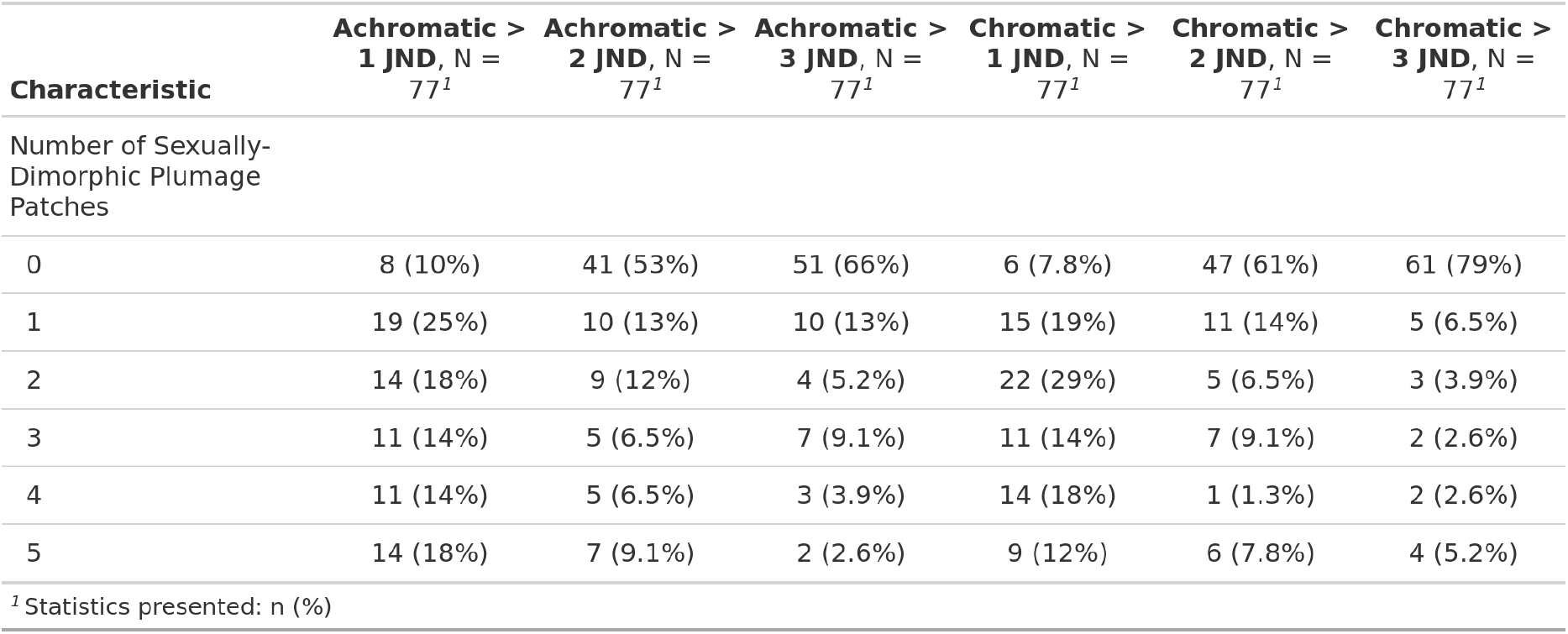
Number of sexually-dimorphic plumage patches for separate achromatic and chromatic just noticeable differences (JND) thresholds by number of *Turdus* thrush species (% of species).

**Fig S1:**
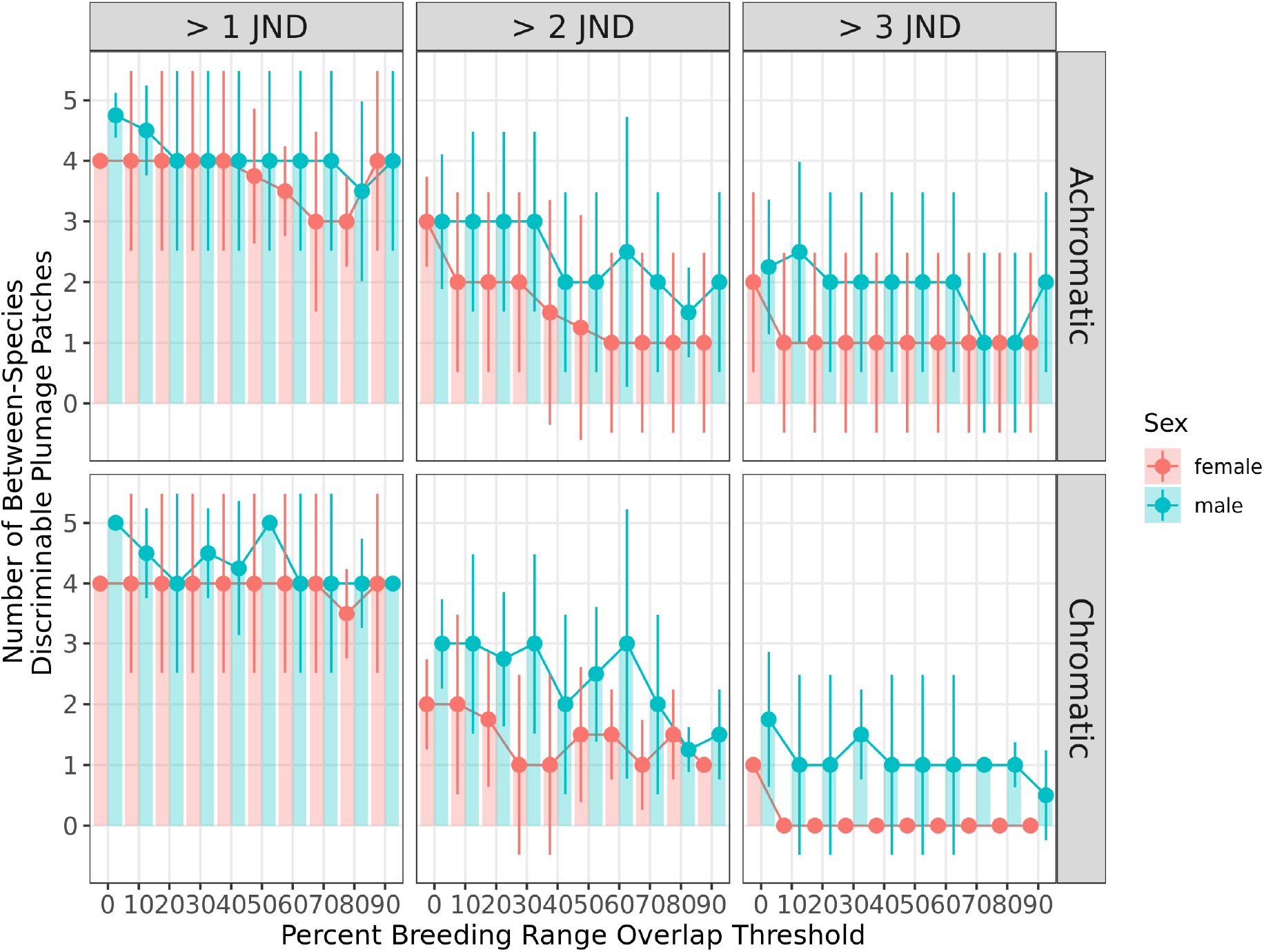
Median ± median absolute deviation of number of distinguishable plumage patches by just noticeable differences (JND) thresholds of 1, 2, and 3 between male and female *Turdus* thrush species in sympatry at various breeding range overlaps (percent).

**Fig S2:**
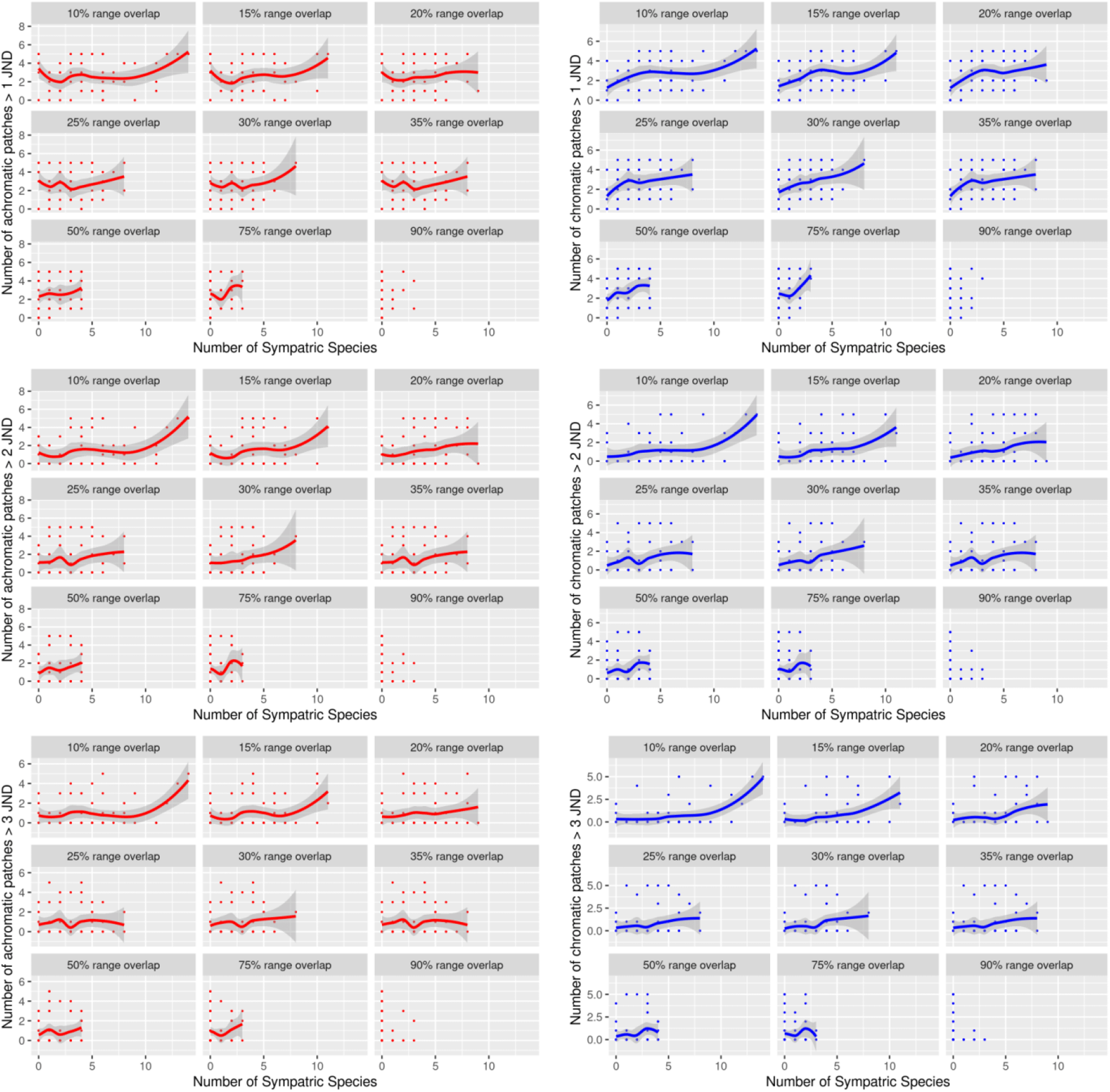
Number of sexually-dichromatic chromatic and achromatic plumage patches versus number of sympatric *Turdus* species, faceted by sympatry overlap thresholds (0-90%). Lines are Loess nonlinear regression fits with no correction for phylogenetic relatedness among species.

**Fig S3:**
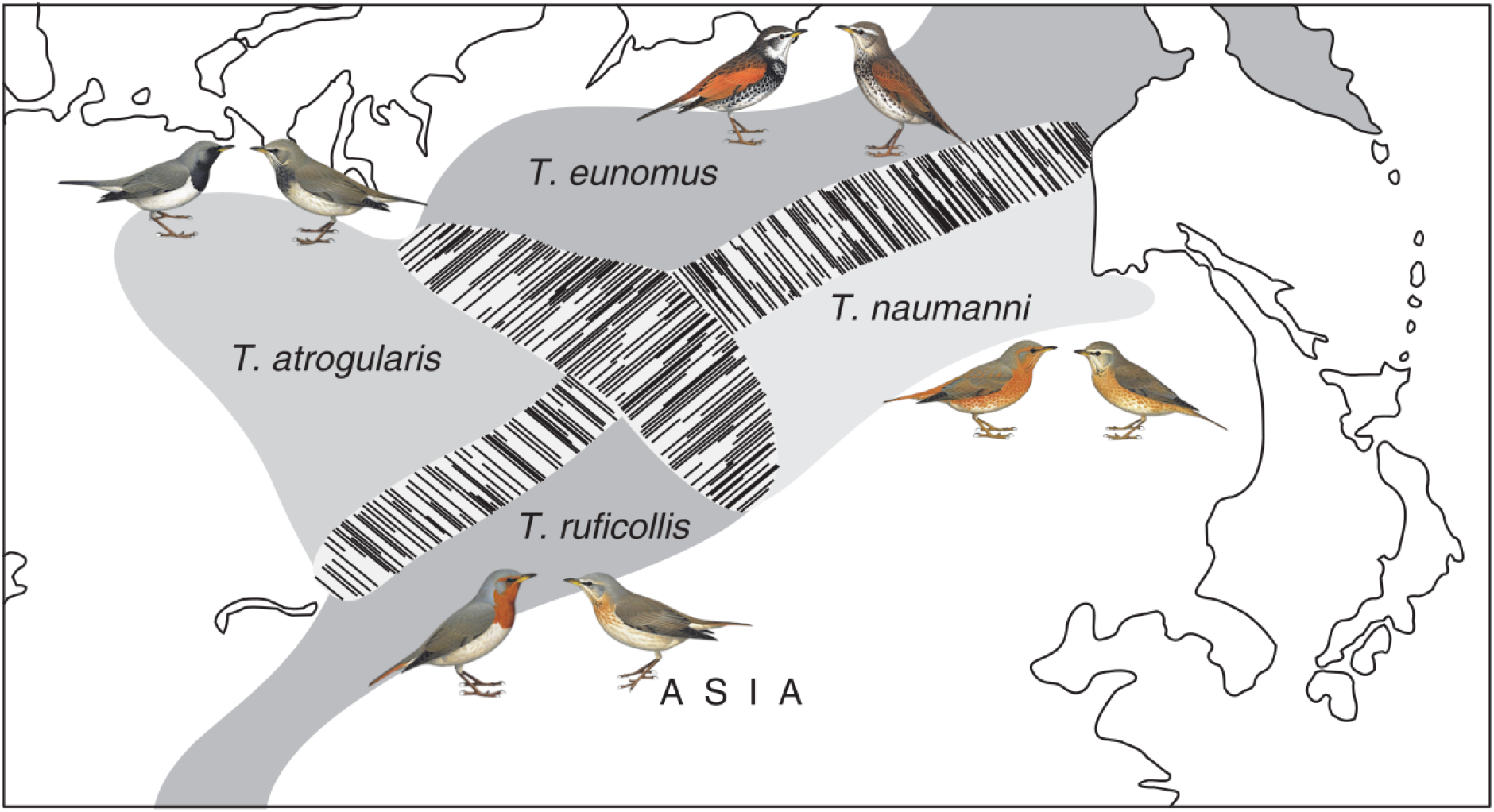
Four species hybrid zone in north-central Asia (*T.atrogularis*, *T.ruficollis*,*T.eunomus*, and *T.naumanni*). Map is from [1]. Illustrations © HBW Alive/Lynx Edicions.

